# Predicting and quantifying coexistence outcomes between resident and invading species using trait and abundance data

**DOI:** 10.1101/2023.01.12.523647

**Authors:** Jocelyn E. Behm, Jacintha Ellers, Wendy A. M. Jesse, Tyler J. Tran, Matthew R. Helmus

## Abstract

A major challenge in invasion ecology is determining which introduced species pose a threat to resident species through competitive displacement. Here, we provide a statistical framework rooted in coexistence theory to calculate coexistence outcomes – including competitive displacement – between resident and invading species. Advantageously, our framework uses readily available trait and abundance data rather than the demographic data traditionally used in coexistence theory applications which is often difficult to collect for most species. Our framework provides methods for *predicting* displacement that has yet to manifest in incipient invasions, and for *quantifying* displacement in ongoing invasions. We apply this framework to the native and introduced gecko species on Curaçao and predict the displacement of all three native species by introduced species and quantify that the displacement of one native species is already underway. Our results affirm that trait and abundance data are suitable proxies to reasonably predict and quantify coexistence outcomes.

## INTRODUCTION

Although invasive species are a major threat to biodiversity (Pyšek *et al.* 2020), not every introduced species negatively impacts resident species and becomes invasive. Given the accelerating spread of introduced species globally (Seebens *et al.* 2018), two major challenges are: 1) *predicting* negative interactions that may happen between introduced and resident species and 2) *quantifying* negative interactions already in progress. This is especially difficult for negative impacts resulting from competitive interactions like displacement (Sheppard 2019). Displacement occurs when an introduced species appropriates a significant portion of, or the entire, spatio-temporal niche of a resident species and is often the first step toward localized extirpation and subsequent extinction of resident species (Gao & Reitz 2017). Despite its importance to invasion ecology, a widely applicable statistical framework for predicting and quantifying displacement is lacking.

Initially developed to understand the assembly of natural communities, modern coexistence theory provides a valuable foundation for predicting and quantifying competitive displacement of resident species by introduced invaders. Under coexistence theory, three coexistence outcomes through competitive interactions are possible: 1) exclusion of the resident through displacement by the invader; 2) exclusion of the invader through biotic resistance from the resident; or 3) coexistence of the resident and invader (Levine *et al.* 2004; Blackburn *et al.* 2014). These coexistence outcomes result from on niche and fitness differences between the competing invader and resident species (Chesson 2000b; MacDougall *et al.* 2009; Kraft *et al.* 2015). Niche differences, like exploiting a unique resource, favor coexistence by reducing interspecific competition and give the invader a population growth rate advantage when the invader’s population size is low, such as when it is first invading. When niche differences are minimal, fitness differences that favor invaders result in the displacement of the resident species (MacDougall et al. 2009, Fig. 1a). In coexistence theory parlance, “fitness differences” represent differences in species’ competitive ability, such as the ability to better capture a shared resource, and give the invader a population growth advantage regardless of its population size (Chesson 2000b).

**Figure 1:**
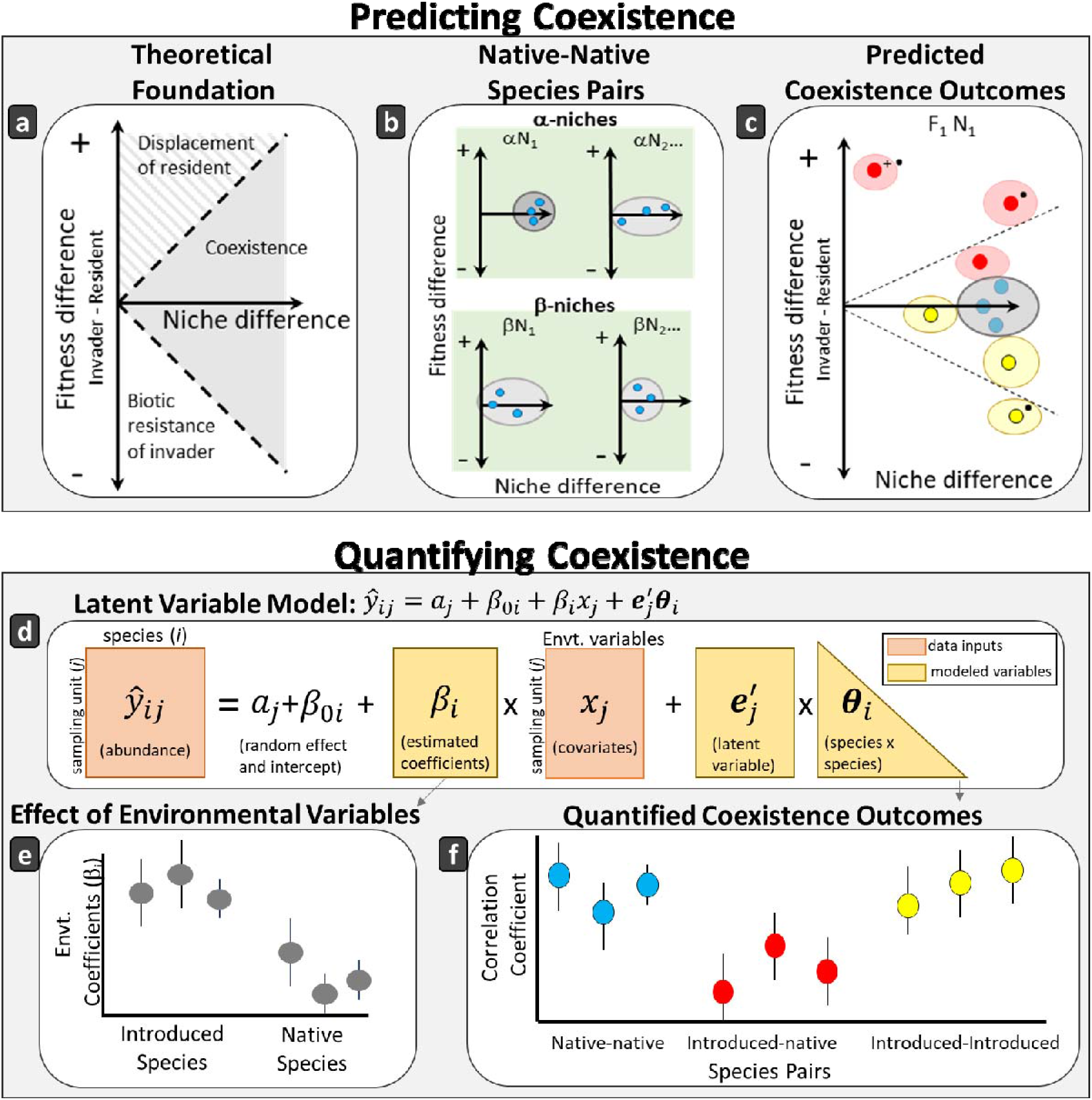
Coexistence outcome framework for predicting and quantifying competitive displacement, biotic resistance, and coexistence among introduced and native species. a) Coexistence theory (modified from MacDougall et al. 2009) for predicting coexistence outcomes is based on fitness and niche differences between invaders and residents. Solid gray area represents coexistence space; the area beyond the dashed lines is exclusion space. The invader is predicted to displace the resident when the invader’s fitness is greater than the resident’s when niche differences are low (striped area), whereas biotic resistance is predicted when the resident’s fitness exceeds the invader when niche differences are low (unshaded area). b) The framework uses niche and fitness differences from native-native species pairs (blue points) to delineate coexistence space as determined by the confidence ellipse of bootstrapped niche and fitness differences among natives (gray ellipses). The α-niche with the greatest niche differences is selected for predicting coexistence outcomes (darkest gray ellipse). Overlap along the β-niche is used to verify that native species pairs co-occur at the spatial scales where competitive exclusion occurs. c) Coexistence outcomes for introduced-native (red points) and introduced-introduced (yellow points) species pairs are determined by the overlap between their confidence ellipse and the native-native pairs’ ellipse. The coexistence space delineated by the dashed lines is determined by the boundary of the confidence ellipses of coexisting species pairs. Species pairs may differ from coexistence space along the fitness (•) and/or niche (+) axes. d) Latent variable models are used for quantifying coexistence outcomes in progress because they account for e) the effect of confounding environmental variables on species abundances and f) provide estimates of correlations between the abundances of pairs of species. Negative correlations indicate exclusion is likely whereas positive correlations support coexistence.

Despite its apparent utility, applications of modern coexistence theory to predicting and quantifying coexistence outcomes are limited to species for which estimating population growth rates through experimental or long-term demographic studies are tractable (Godoy & Levine 2014; Ocampo-Ariza *et al.* 2018; Sheppard 2019). While these studies have greatly advanced coexistence theory, methods reliant on population growth rate data are not applicable for many species on Earth. A key finding from these studies, however, indicates that other types of data, such as species traits and population abundances, can be used to infer coexistence outcomes (Adler *et al.* 2013; Kraft *et al.* 2015; Letten *et al.* 2017).

Here, we make coexistence theory applicable to a wider range of species by providing a statistical framework for calculating coexistence outcomes based on proxies for growth rate data that are logistically feasible to measure, especially during incipient invasions when data are scarce. For *quantifying* the three coexistence outcomes – coexistence, biotic resistance, or displacement – occurring between resident and invading species pairs, we use the relative abundances of species across sites as proxies for differences in population growth rates (Blanchet *et al.* 2020). Although data intensive, quantifying coexistence outcomes provides a direct estimate of competitive exclusion in progress. In the absence of widespread abundance data, we use species-level trait and habitat use data to *predict* coexistence outcomes. In particular, we use traits associated with resource partitioning and competitive advantages as proxies for niche and fitness differences, respectively (MacDougall *et al.* 2009). We apply our framework to a case study of three native and three introduced gecko species on Curaçao. While our framework is illustrated using geckos, we provide general guidelines so that it can be applied to other systems for which trait or abundance data are available.

## COEXISTENCE OUTCOME FRAMEWORK

### Predicting pairwise coexistence outcomes with trait data

As a theoretical foundation, we use MacDougall et al. (2009) for predicting three coexistence outcomes – coexistence, biotic resistance, or displacement – between invader-resident species pairs based on niche and fitness differences (Fig. 1a). Using invasibility model terminology (Chesson 2000b), ‘resident’ is the already-established species and ‘invader’ is the species that is establishing, therefore residents can be native or introduced species. Although we calculate all pairwise species coexistence outcomes, we highlight displacement of native species due to conservation concerns.

#### Niche differences

Coexistence theory suggests that traits related to local-scale resource partitioning, known as a species’ α-niche, are involved in stabilizing coexistence mechanisms (Silvertown 2004; Ackerly *et al.* 2006). When overlap in these traits occurs among species, local-scale competitive exclusion is probable. In comparison, a species’ β-niche represents its broader-scale habitat tolerances. Differences in β-niche traits facilitate broader-scale coexistence for species that overlap in α-niche traits by reducing their spatial and/or temporal overlap (Chesson 1985, 2000a; Gross *et al.* 2013).

In our framework, we calculate differences in the α-niche for predicting coexistence outcomes. Differences in the β-niche are also calculated to assess the spatiotemporal overlap of species because species with high β-niche differences likely do not interact at the scales where exclusion is possible. We assert that α- and β-niches are best estimated from species’ trait and habitat use data collected in the invaded range, as traits and habitat use could differ for species between their invaded and native ranges (e.g., Kolbe *et al.* 2012). However, for some invaders, especially in the early stages of invasion when populations are localized, the totality of the invader’s β-niche may not be realized, providing illusory broad-scale habitat partitioning.

To calculate niche differences, we use niche overlap metrics that can handle the myriad formats that trait data can take (e.g., discrete, binary, continuous) (Mouillot *et al.* 2005; Geange *et al.* 2011). These methods calculate niche overlap by generating nonparametric curves that represent the probability density functions of the niche data using either kernel density estimation or mixture model methods (see Mouillot *et al.* 2005). This avoids the assumption of normally distributed (or other distribution) data and niche overlap is calculated as the overlapping area between the probability distributions (Geange *et al.* 2011). Although niche overlap is a metric of niche similarity, it can easily be converted to niche differences and used in our framework.

#### Fitness differences

Under coexistence theory, fitness differences describe variation in population growth rates among species due to differences in competitive ability over shared resources (Chesson 2000b; Adler *et al.* 2013). In our framework, to represent fitness differences, traits can be used that capture differences in competitive dominance among species such as traits associated with exploiting limiting resources, variation in predator or pathogen susceptibility, and population-level differences in fecundity or growth rate (MacDougall *et al.* 2009). Depending on the system, population-level variables that approximate growth rates, such as reproductive output, may better capture fitness differences and be more logistically tractable to measure than specific traits. To calculate fitness differences, we advocate using multiple, standardized traits recorded from the invaded range and distance methods to calculate differences among them, rather than single traits, when available.

#### Coexistence space

Under coexistence theory, particular combinations of niche and fitness differences generate a ‘coexistence space’ and pairs of species with niche and fitness differences that fall within this space should coexist (MacDougall *et al.* 2009) (Fig. 1a). Traditional coexistence theory applications define coexistence space using demographic data derived experimentally or from long-term studies to estimate population growth rates (Godoy & Levine 2014; Ocampo-Ariza *et al.* 2018; Sheppard 2019). A defining feature of our framework is that we delineate coexistence space using coexisting native species pairs. We select co-occurring native species that are presumed to coexist and confirm their coexistence based on their niche and fitness differences (Siepielski & McPeek 2010).

To delineate coexistence space, we identify the traits associated with the maximal α-niche and minimal fitness differences that underlie coexistence in the native species pairs, under the assumption that these are the traits most strongly associated with coexistence in the system (Fig. 1b). Once these traits are identified, we use the confidence ellipse that bounds bootstrapped values of all native-native species pairs’ niche and fitness differences calculated from these traits (Ocampo-Ariza *et al.* 2018) for delineating the coexistence space boundary that marks the transition between coexistence and exclusion. We also confirm that native species with high α-niche differences have low β-niche differences indicating that they are coexisting at local spatiotemporal scales (Siepielski & McPeek 2010). While we advocate using native-native species pairs given their likelihood for coexistence, using introduced residents in pairs may be suitable, assuming that sufficient time has passed for the resident to have naturalized and coexist with native species (e.g., Heard & Sax 2013). Using recently introduced residents may yield misleading results as exclusion may still be underway.

#### Coexistence outcome

Under coexistence theory, the species pairs with niche and fitness differences that fall outside coexistence space should experience competitive exclusion. In our framework, the coexistence outcomes of invader-resident species pairs are determined by the location of their niche and fitness differences relative to the coexistence space delimited by the native-native species pairs. For invader-resident pairs with predicted exclusion outcomes, we identify the directionality of the exclusion – displacement or biotic resistance – and whether it falls outside the coexistence space along the niche and/or fitness axes (Fig. 1c). To determine the coexistence outcomes for invader-resident pairs, we calculate the confidence ellipses of bootstrapped niche and fitness differences for each invader-resident pair. Invader-resident pair confidence ellipses that do not overlap with the native-native species confidence ellipses have a predicted exclusion outcome. The full boundaries of the coexistence space, then, are determined by the niche and fitness differences of the coexisting invader-resident and native-native pairs (Fig. 1c).

### Quantifying coexistence outcomes with abundance data

Under coexistence theory, the abundances of invader and resident species pairs that are currently experiencing competitive exclusion should negatively covary (Ulrich *et al.* 2017; Zurell *et al.* 2018). However, environmental variables that affect invading and resident species differently may also cause negative covariance among abundances in the absence of competitive interactions between invaders and residents. For example, a survey along an environmental gradient may reveal a negative covariance in the abundance of residents and invaders due to differing tolerances to the environmental variable rather than due to negative species interactions. Therefore, to statistically quantify coexistence outcomes, models are needed that can estimate the directionality of covariances between the abundances of invader and resident species while accounting for potentially confounding environmental variables (O’Reilly-Nugent *et al.* 2020).

In our framework, we use joint species distribution latent variable models (LVM) to estimate correlations in species abundances after accounting for environmental variables (for a full statistical overview see Pollock *et al.* 2014; Warton *et al.* 2015; Ovaskainen *et al.* 2017). In general, LVMs are similar in structure to hierarchical generalized linear mixed models, however species-species correlations are modeled through a random effect term at the level of the sampling unit that includes measured and unmeasured (‘latent’) predictors (Warton *et al.* 2015) (Fig. 1d). The loadings of these latent predictors provide an indication of whether a pair of species co-occur more or less often than expected by chance after accounting for the influence of environmental variables on species abundances (Ovaskainen *et al.* 2017) (Fig. 1e). Negative correlations between resident-invader species pairs in LVMs provide support for competitive exclusion (Warton *et al.* 2015).

To calculate coexistence outcomes, we use LVMs to estimate pairwise correlations in species abundances to provide an indication of coexistence outcomes in progress (Fig. 1f). We fit LVMs using environmental data and species abundances collected during observational field studies at the predicted scale that species interactions are likely to occur (Zurell *et al.* 2018). We strongly urge the use of abundance data over presence/absence data because negative biotic interactions are likely manifest as shifts in population abundances before outright extirpation occurs (Radinger *et al.* 2019; Blanchet *et al.* 2020). While LVMs, like any statistical modelling approach of this nature, are correlative and do not identify a definitive mechanism for a coexistence outcome, they provide an indication that a concerning coexistence outcome like displacement could be occurring and form the basis for developing experimental hypotheses (Ovaskainen *et al.* 2017).

## CASE STUDY

### Study system and data collection

We applied our framework to three native and three introduced gecko species from Curaçao, the largest island (444km^2^; 12.187°N, −68.983°W) in the Caribbean Leeward Antilles ca. 70km north of Venezuela (Fig. 2). The three native gecko species on Curaçao, *Thecadactylus rapicauda, Phyllodactylus martini*, and *Gonatodes antillensis*, have island-wide distributions and are often found co-occurring at the same sites. *Hemidactylus mabouia* was the first gecko introduced to Curaçao in the late 1980s and now has a widespread distribution predominantly in developed habitats (van Buurt 2011). Anecdotal evidence suggests *H. mabouia* is displacing *P. martini* and possibly *G. antillensis* (Dornburg *et al.* 2011, 2016; van Buurt 2011), and an observational study supports displacement based on a negative covariance between *H. mabouia* and *P. martini* abundances, yet the possible confounding effects of environmental variables were not accounted for (Hughes *et al.* 2015). Two additional geckos, *Lepidodactylus lugubris* and *H. frenatus*, are at the incipient stages of their invasion, likely introduced to Curaçao since 2000 (Behm *et al.* 2019). A fourth gecko, *Gekko gecko*, was more recently introduced, but was excluded from this study because it is likely a predator and not a competitor of the other six species (Behm *et al.* 2019; Perella & Behm 2020).

**Figure 2:**
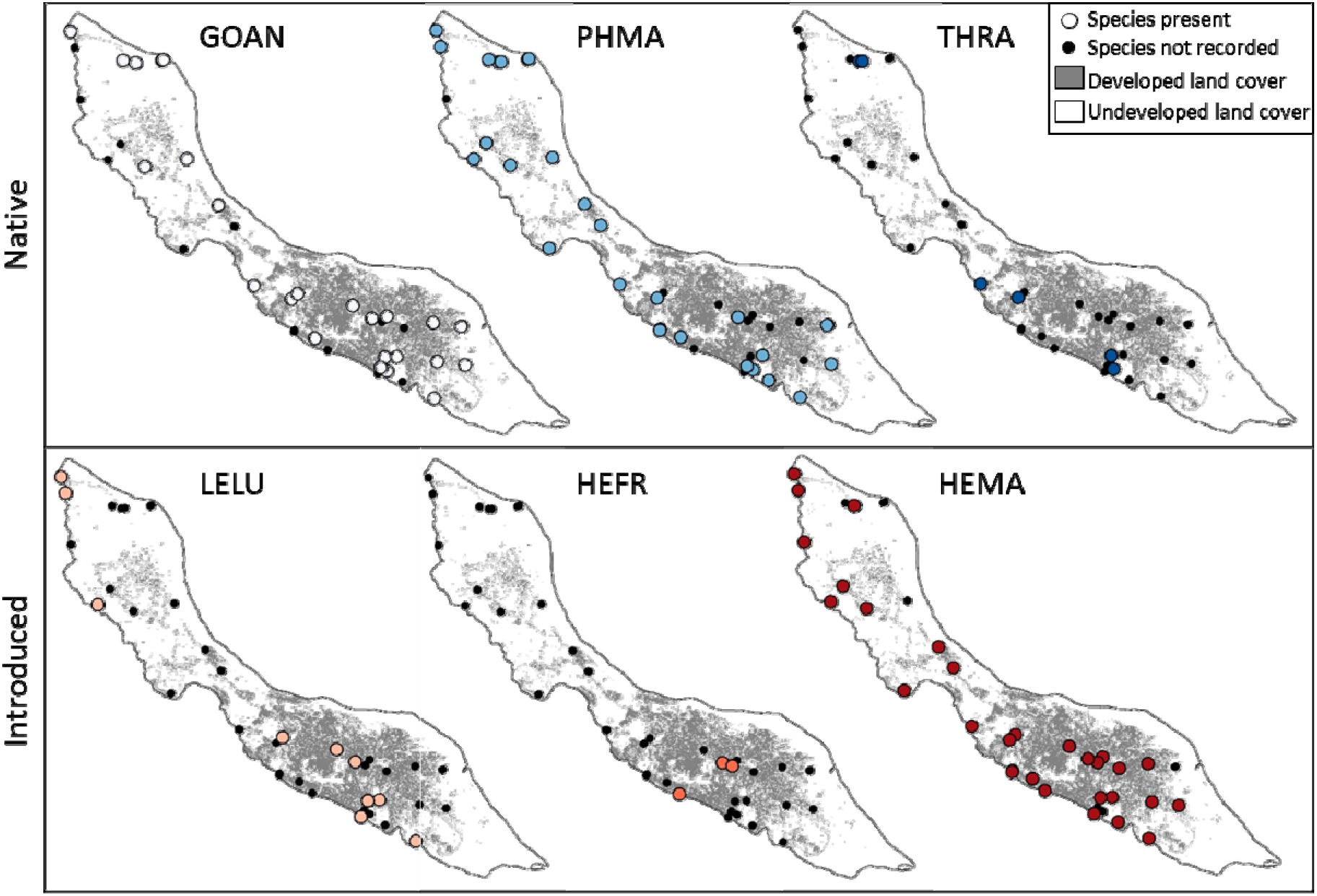
Locations of native (top) and introduced (bottom) gecko species found on Curaçao island across 41 survey sites spanning a gradient of development in the surrounding landscape. Species name abbreviations are as follows: *Gonatodes antillensis* – GOAN; *Phyllodactylus martini* – PHMA; *Thecadactylus rapicauda* – THRA; *Lepidodactylus lugubris* – LELU; *Hemidactylus frenatus* – HEFR; *Hemidactylus mabouia* – HEMA. Native species are shown in shades of blue and introduced species are shown in shades of red.

We conducted gecko surveys at sites across Curaçao from January 27 – March 11, 2017 at the start of the rainy season when all six species are reproducing. Based on our work on Curaçao and similar islands (Jesse *et al.* 2018; Behm *et al.* 2022), we anticipated habitat development would be a strongly influential environmental variable that the native and introduced gecko species would respond to differently and therefore structured our sampling to span the gradient of habitat development on Curaçao. Surveys were conducted between one and four hours after dusk, which corresponds to peak gecko activity, at “undeveloped” (>100m from buildings and/or paved surfaces), and “developed” (<20m from buildings and/or paved surfaces) sites. Undeveloped sites were generally nature preserves and hiking areas, while developed sites were mostly residential yards or resort gardens. To quantify gecko relative abundances to use in LVMs, we used an area-constrained approach such that the total area searched at each site was approximately 400m^2^ and all gecko species per age class (adult or juvenile) were recorded. We also recorded microhabitat use of individuals and captured a subset of the individuals to record snout-vent-length (SVL) and take standard photos for measuring additional morphological traits and reproductive status for quantifying α-niche and fitness metrics, respectively. To understand the full distribution of geckos in habitats across the island for quantifying β-niche, we conducted additional area-unconstrained searches which consisted of searching a haphazardly established transect and recording the geckos encountered. In total, data from 288 geckos across 41 sites were used for our α-niche, β-niche and fitness difference calculations, and abundances of 837 individuals recorded during area-constrained surveys at 26 sites were used for LVM analyses (Table S1).

### Niche Differences

To represent the α-niche, we used standard morphological traits and microhabitat use variables associated with resource use and competitive interactions in squamates (Beuttell & Losos 1999; Mahler *et al.* 2013). We recorded the following morphological traits from the photos taken in the field using ImageJ software (imageJ.nih.gov/ij): snout-vent-length (SVL), snout width (SW), head width (HW), inter-eye-distance (IED), finger 4 width (F4W), finger 4 length (F4L), finger 4 lamellae count (FLC), toe 4 length (T4L), toe 4 width (T4W), and toe 4 lamellae count (TLC). During surveys, we also collected the following microhabitat use data for each individual: perch height (cm) (continuous), perch diameter (cm) (continuous; flat perches were assigned maximum perch diameter), perch lighting (binary: light / dark), and perch substrate (binary: natural/manmade). All continuous α-niche traits were log-transformed and scaled to have unit variance prior to analyses. Because many of these α-niche traits were significantly correlated (Fig. S1), we eliminated the collinearity effects by conducting a principal components analysis (PCA) on all α-niche variables and retained the first two axes that explained >70% of the variation for subsequent niche overlap analyses (hereafter PC1α, PC2α). Traits and habitat use variables that scale with gecko body size (body size, head dimensions, perch height and digit length) loaded positively on PC1α whereas microhabitat partitioning traits (digit lamellae counts) and habitat use variables (perch diameter, perch lighting, perch substrate) loaded positively and negatively onto PC2α, respectively (Table S2). We interpret PC1α as a resource use axis (i.e., trophic resources), and PC2α as a microhabitat use axis.

To quantify β-niche, we performed a supervised land cover classification at a 30 m resolution to estimate the broad-scale habitat types on Curaçao (Appendix S1A). From this classification, the proportion of developed habitat, low-density, and high-density vegetation were estimated within 100, 250, 500, 750, and 1000 m radius buffers around each survey site. Due to significant correlations among these three β-niche land-cover variables (Fig. S2), we applied PCA analysis and retained the first two axes that explained >80% of the variation (hereafter PC1β, PC2β) for subsequent analyses. Developed habitat loaded positively onto PC1β, whereas low-density vegetation loaded positively, and high-density vegetation loaded negatively onto PC2β (Table S3). We interpret PC1β as a development axis and PC2β as a vegetation structure axis.

We quantified pairwise niche overlap along both α-niche (PC1α, PC2α) and β-niche (PC1β, PC2β) dimensions, setting all four variables to continuous (Geange *et al.* 2011). We calculated niche differences to use in coexistence outcome prediction analyses by subtracting niche overlap from one. To distinguish the directionality of biotic resistance versus displacement exclusion outcomes (Fig. 1a), we designated the species within pairs that had been on Curaçao the longest as the resident and for native-native species pairs, one of the species was randomly designated as the resident.

### Fitness Differences

To quantify fitness differences, we used three population-level variables measured during our field surveys that approximate population growth rates and integrate differences in competitive ability: mean clutch size per female, proportion of gravid females, and proportion of juveniles. These variables are directly related to population-level differences in fecundity (MacDougall *et al.* 2009) and are the most logistically tractable variables associated with fitness differences that can be measured in our system during an incipient invasion.

For all species except *T. rapicauda*, eggs are identifiable through the translucent abdominal skin of gravid females. From the photos taken in the field, we calculated the proportion of gravid females (out of all females) during area-constrained surveys, and recorded clutch sizes. For *T. rapicauda*, we could not quantify the proportion of gravid females, but we did include the average clutch size for *T. rapicauda* reported in the literature (Vitt & Zani 1997). The proportion of juveniles was quantified from area-constrained survey data. We confirmed that the proportions of juveniles and females were invariant with respect to sampling date and habitat type (all *P*>0.05, Table S4, S5).

To quantify pairwise fitness differences to use in subsequent coexistence outcome analyses, we calculated Euclidian distances among metrics using the dist function in R on the matrix of species-level means of our three fitness metrics.

### Coexistence Space and Coexistence Outcomes

To delineate the coexistence space for calculating coexistence outcomes (Fig. 1a), we plotted each of the four niche difference dimensions (PC1α, PC2α, PC1β, PC2β) against our single fitness difference metric and identified the α-niche dimension that native-native pairs partitioned most strongly and visually confirmed low niche differences for native-native pairs along β-niche dimensions (Fig. 1b, Fig. S3). We used this PC1α niche dimension along with our fitness difference values to calculate coexistence outcomes among all species pairs (Fig. 1c).

### Quantifying Coexistence Outcomes

To determine whether covariance patterns in the abundances of introduced-native species pairs are due to biotic interactions rather than habitat development, we separated the effects of development and species interactions by modeling gecko abundances from area-constrained surveys using an LVM with a Poisson distribution (Fig. 1d). We modeled the observed species abundance data *y_ij_* as:

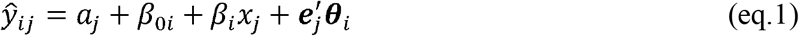

where 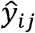 is the predicted abundance of gecko species *i* at site *j.* The random effect *a_j_* accounts for differences among sites in total assemblage abundance and the intercept *β*_0*i*_ accounts for differences in the overall abundance of each species across all sites (Ives & Helmus 2011). The effect of habitat development on each species is estimated in *β_i_x_j_*, where *β_t_* is a fixed slope for each species and *x_j_* is habitat development, standardized to have mean=0 and standard deviation=1. The random term 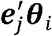 consists of two latent variables stored in 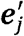, and species responses to those variables as the latent-variable regression coefficients ***θ**_i_*. Interspecific interactions are inferred as the pairwise correlations *ρ_ii_* from the covariance of 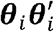 (Warton *et al.* 2015).

Despite restricted distributions of *T. rapicauda* and *H. frenatus* at sites (Fig. 2, Table S1), we included them in LVMs, with a cautious interpretation of their results. We fit separate LVMs to each of our five habitat development buffer radii generated for our β-niche analysis to explore the response to development and species interactions using the gllvm R package (Niku *et al.* 2019; see Appendix S1B for selecting LVM R packages). Significance of the development effects, *β_i_x_j_*, were assessed with 95% confidence intervals (Fig. 1e) and the significance of the interspecific correlations (Fig. 1f) was assessed with likelihood ratio tests of the full model (eq. 1) versus the null model without 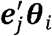 (Niku *et al.* 2019).

## RESULTS

### Niche and Fitness Differences

The first two axes of a principal components analysis (PCA) explained 72.41% of the variation in α-niche traits (resource use axis - PC1α: 59.42%; microhabitat axis - PC2α: 12.99%). The three native species did not overlap on PC1α, yet native *P. martini* did overlap with all three introduced species on PC1α (Fig. 3a). For the β-niche, the first two axes of a PCA explained 85.45% of the variation in β-niche habitat use traits (development axis - PC1β: 56.35%, vegetation structure axis - PC2β: 28.10%). While native species were skewed more towards the negative end of PC1β (lower development), there was much higher overlap among all six species in β-niche compared to α-niche traits (Fig. 3b). These patterns were confirmed by the distribution of species across the island; introduced species were found predominantly at developed sites whereas native species were found at both developed and undeveloped sites (Fig. 2).

**Figure 3:**
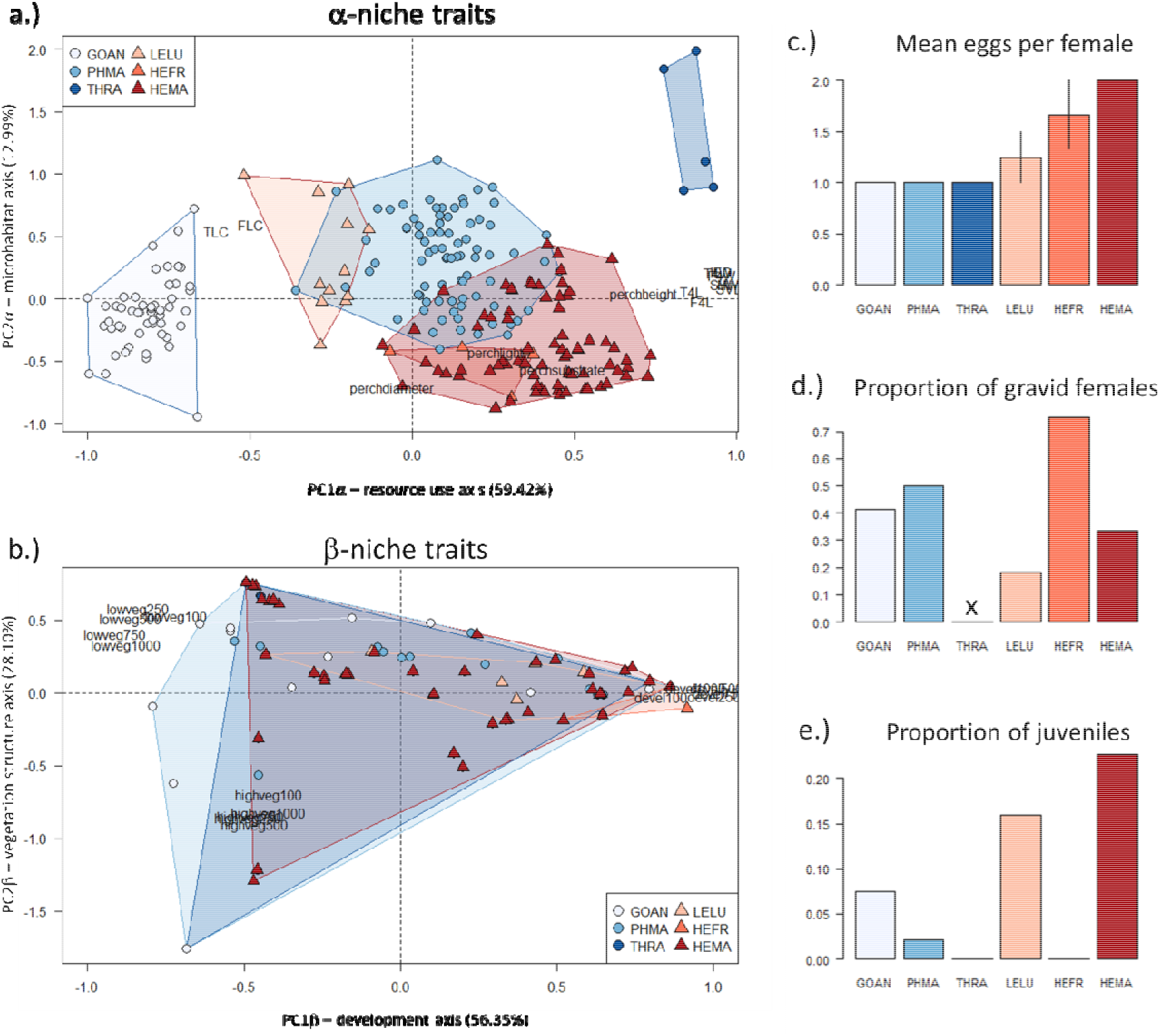
Niche and fitness traits used in coexistence outcome calculations. We conducted a principal components analysis (PCA) on α-niche trait data (a), and broad-scale habitat use β-niche data (b) (see text for variable abbreviations in PCA plots) collected from individuals of the six species we surveyed on Curaçao island. We used three metrics in our fitness difference calculations: mean number of eggs per gravid female (mean +/− SE) (c), proportion of gravid females (X indicates data are not available for that species) (d), and proportion of juveniles (e). Native species are shown in shades of blue and introduced species are shown in shades of red. Species name abbreviations are given in Fig. 2.

The native and introduced species differed in our three species-level fitness metrics: mean number of eggs per female, proportion of gravid females, and proportion of juveniles (Fig. 3c-e). The native species, *G. antillensis*, and *P. martini*, consistently had only one egg per gravid female recorded from field measurements as did *T. rapicauda* as reported from the literature. In comparison, the introduced species varied between one or two eggs (*L. lugubris* and *H. frenatus*) or were invariant with two eggs per gravid female (*H. mabouia*) (Fig. 3c). Apart from *H. frenatus*, the introduced species had a lower proportion of gravid females (Fig. 3d) and a higher proportion of juveniles (Fig. 3e) than the native species.

### Predicting Coexistence Outcomes

To define coexistence space, the comparison of our two α-niche metrics showed that the native-native species pairs exhibited higher niche differences on the resource use axis, PC1α, than the microhabitat use axis, PC2α (Fig. S3). We confirmed that the native species pairs showed lower niche differences on β-niches than α-niches (Fig. S3) suggesting that the native species are coexisting at local spatial scales by partitioning trophic resources (mean niche differences for native-native species pairs: PC1α: 1.00; PC2α: 0.71; PC1β: 0.42; PC2β: 0.38). Therefore, we used PC1α to define coexistence space in subsequent coexistence outcome analyses.

Our prediction of coexistence outcomes revealed that six species pairs, five of which included introduced *H. mabouia*, had predicted exclusion outcomes, and four of these pairs were in the direction of the displacement of native species (Fig 4). Three pairs, *H. mabouia* with *T. rapicauda, G. antillensis* and *L. lugubris*, differed from the coexistence space only along the fitness axis, one pair of introduced species, *H. frenatus-H. mabouia*, differed from coexistence space along the niche axis only, and the pair, *H. mabouia-P. martini* differed from coexistence space along both niche and fitness axes.

**Figure 4:**
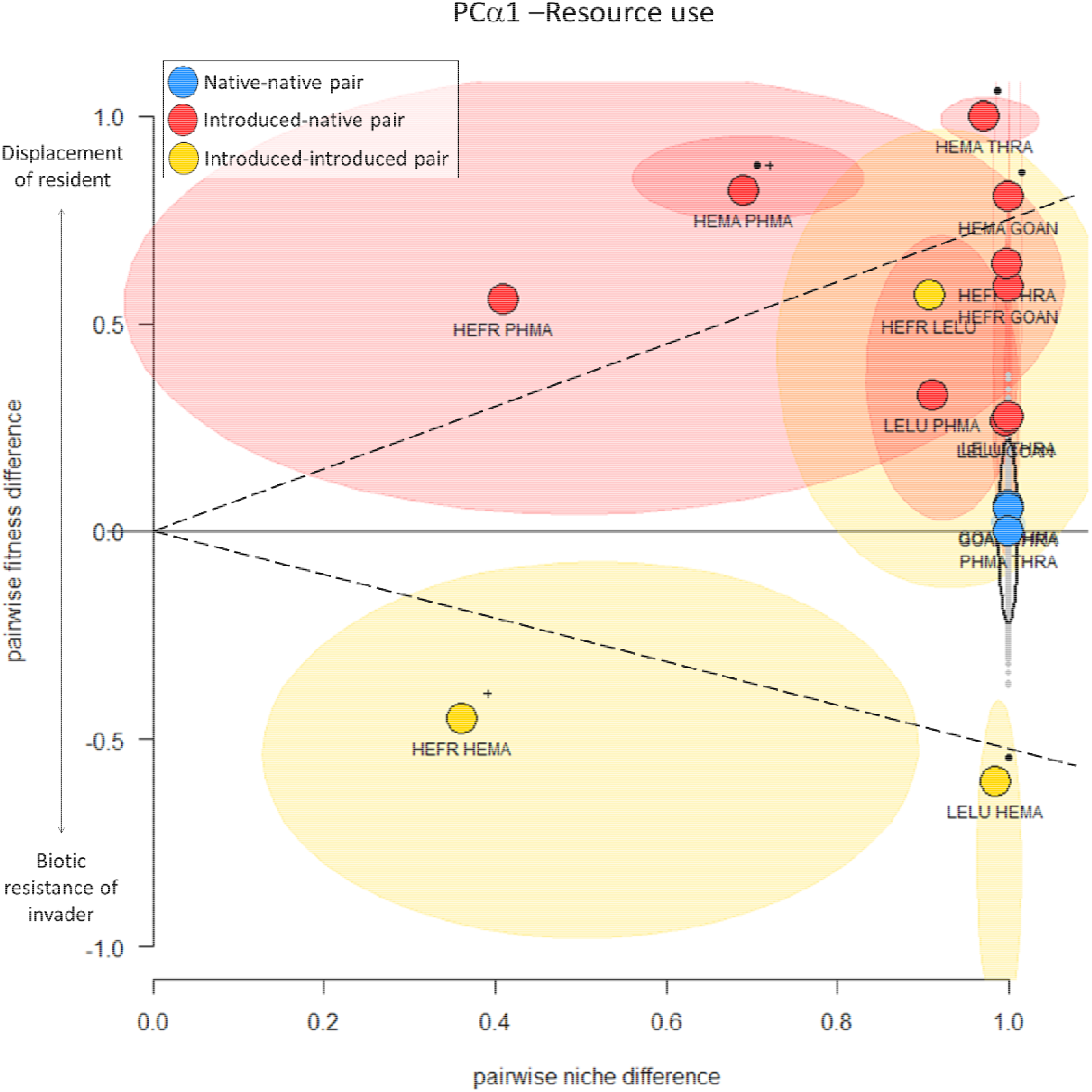
Predicted coexistence outcomes of Curaçao island geckos. Pairwise fitness and niche differences along PC1α (resource alpha) for native-native, native-introduced and introduced-introduced gecko species pairs. The gray point cloud indicates the bootstrapped values for the three native-native species pairs and is bound by the 95% confidence ellipse (black line). Points are labeled with species pair names with the species designated as the invader listed first using same abbreviations as in Fig. 2. Species-pair points are surrounded by their 95% confidence ellipse to show which confidence ellipses intersect the confidence ellipse of the native-native species pairs. Points with a (+) have confidence ellipses that do not overlap with the confidence ellipse along the niche axis, while points with a (•) do not overlap along the fitness axis. The dashed line delineates the boundary of coexistence space as determined by the points with a predicted coexistence outcome.

### Quantifying Coexistence Outcomes

The joint species LVMs indicated a positive effect of habitat development on the abundances of introduced *H. mabouia* and *L. lugubris*, and a negative effect of development on native *P. martini* and these effects were consistent across development buffer sizes (Fig. 5, Fig. S4). Likelihood ratio tests indicated that including interspecific species correlations significantly increased the model fit (all *P*<0.001, Table S6). Overall, native species consistently had positive or neutral correlations when paired with other native species, supporting coexistence (Fig. 6). Similarly, introduced species had positive correlations when paired with other introduced species (Fig. 6, Fig. S5), indicating the negative interactions we predict (Fig. 4) have yet to manifest. However, the two native species, *P. martini* and *T. rapicauda*, consistently had negative correlations when paired with introduced species and the introduced species, *H. mabouia* and *H. frenatus* had negative correlations when paired with native species (Fig. 6, Fig. S5). There was a strong negative correlation between *P. martini* and *H. mabouia* (*ρ*= −0.98) which supports displacement in progress for *P. martini.* Strong negative correlations were also identified between *H. mabouia* and *T. rapicauda* (*ρ*= −0.99), as well as between *H. frenatus* and both *P. martini* (*ρ*= −0.92) and *T. rapicauda* (*ρ*= −0.82), however, we interpret these final two results cautiously due to the low sample sizes of *H. frenatus* and *T. rapicauda.*

**Figure 5:**
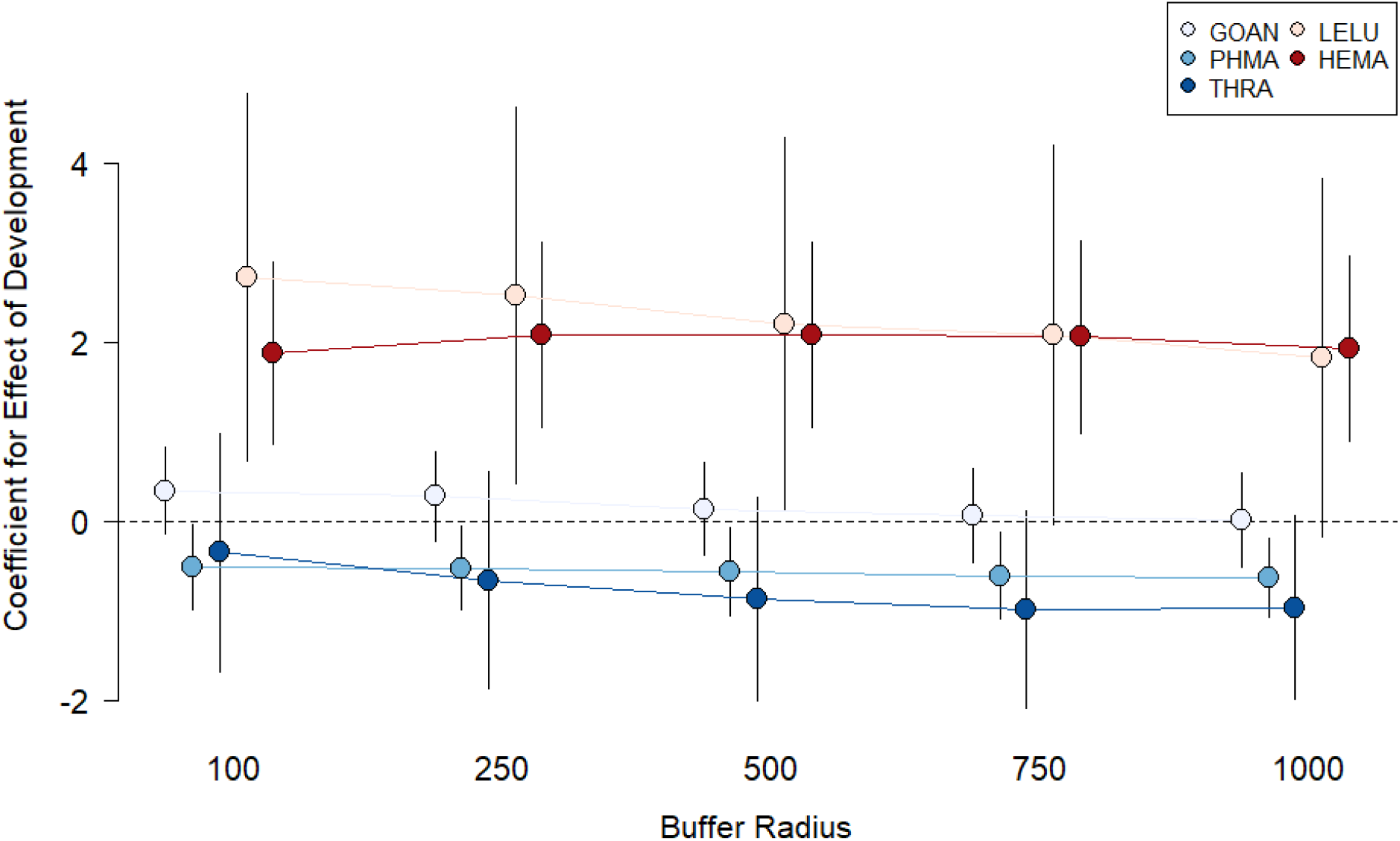
The effect of development (coefficient +/− 95% confidence interval) at 5 spatial scales (buffer radii) on the abundance of five gecko species (HEFR excluded for readability due to high variability, see Fig. S4 for HEFR) as indicated from latent variable model. Native species are in shades of blue and introduced species are in shades of red. Species name abbreviations are listed in Fig. 2.

**Figure 6:**
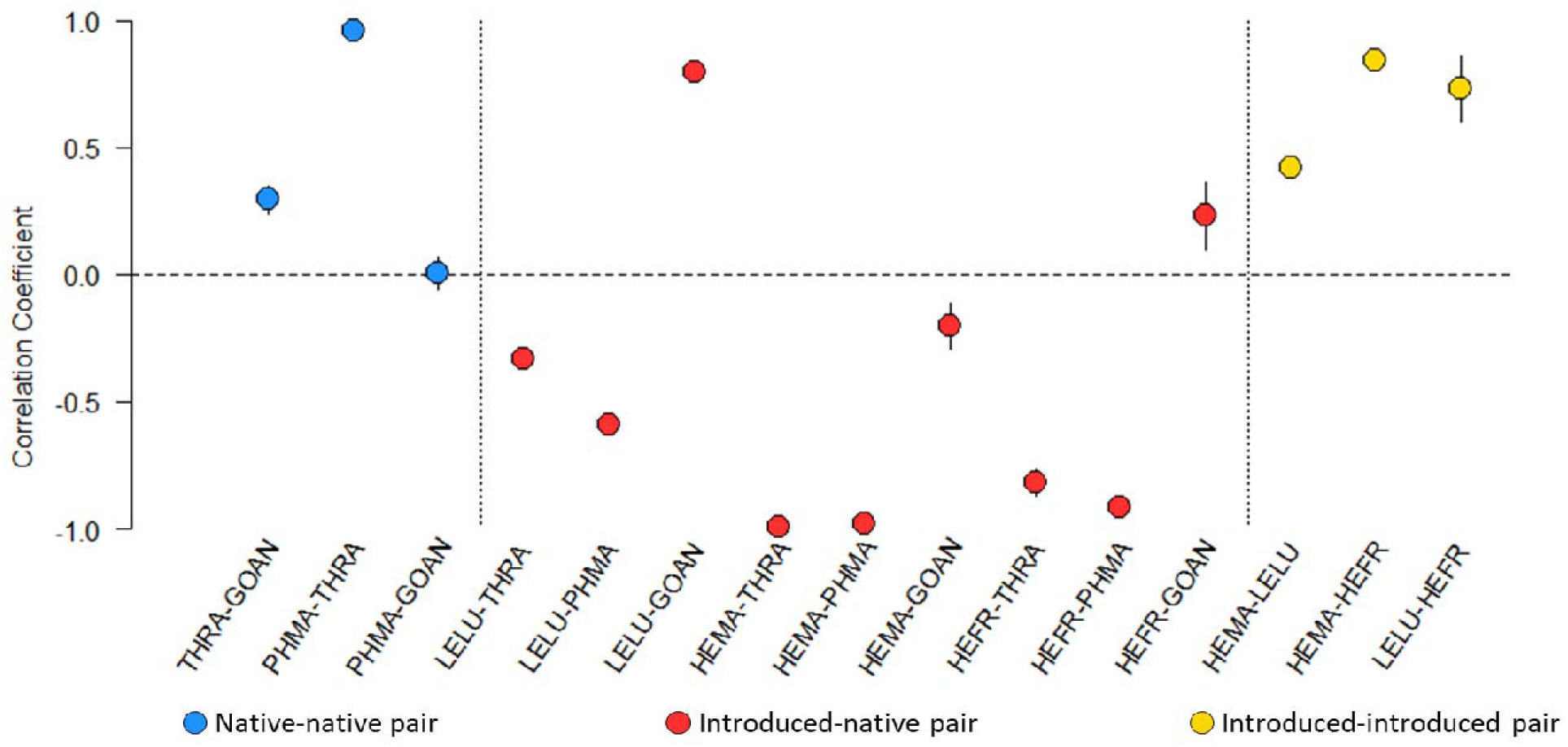
Quantified coexistence outcomes of Curaçao gecko species. Points are mean pairwise interaction correlation coefficients across buffer sizes (+/− standard error) calculated from latent variable models after accounting for the effect of development on each species. Colors of points indicate the type of species pairings. Positive values indicate the species’ abundances across sites are positively correlated whereas negative values indicate negative correlations in abundances across sites and is consistent with competitive displacement. Species name abbreviations are listed in Fig. 2.

## DISCUSSION

Considering that the current rates of species introductions will accelerate, statistical frameworks are needed for calculating the competitive displacement of residents by introduced species. Rooted in coexistence theory, our framework provides a widely applicable method for *predicting* and *quantifying* coexistence outcomes between invading and resident species (Fig. 1). Application of our framework predicted the displacement of all three native gecko species and quantified that the displacement of the native, *P. martini*, is already underway. These results provide useful indications of where conservation and management efforts should be allocated as well as directions for future studies.

The prediction component of the framework can be used in situations where widespread abundance data are lacking due to logistical constraints and/or the early stages of an invasion when species distributions are limited. However, sufficient knowledge of the system is still needed to select candidate traits that underlie niche and fitness differences in the system. Consistent with other native Caribbean squamates, morphological traits best represented interactions underlying coexistence in our island system (Mahler *et al.* 2013). This morphological divergence likely allows Curaçao’s three native gecko species to use the same microhabitats while still partitioning resources. In different systems, coexistence may be maintained through microhabitat partitioning, behavioral differences, or other mechanisms (e.g., Kent & Sherry 2020). For fitness differences, we combined population attributes (proportions of gravid females and juveniles) and traits (number of eggs per female) into a single fitness metric. In other systems, different population-level attributes or traits measured from individuals may be more tractable and using the approach of testing multiple fitness metrics to delineate coexistence space may be appropriate.

The quantification component of our framework requires abundance data collected across enough sites to fit an LVM, making it applicable only to invasions where the species has spread to many locations. In addition, LVMs have the limitation of only providing symmetrical correlation coefficients for species pairs. When displacement is occurring, competition is unlikely to be symmetrical, and the correlation coefficient does not provide an indication of the winner/loser of the interaction. Additional knowledge from the system will be needed to make those inferences (O’Reilly-Nugent *et al.* 2020). Nonetheless, LVMs are powerful tools for inferring species interactions. Furthermore, although the prediction and quantification components of the framework can be used independently, using them in a complementary fashion provides a holistic view of the study system. Specifically, the predicted results can help with the interpretation of the LVM correlations, whereas the LVM correlations can confirm the predicted coexistence outcomes.

The application of our framework to the geckos on Curaçao supports the claims *H. mabouia* is displacing *P. martini* (van Buurt 2005, 2011; Dornburg *et al.* 2011, 2016; Hughes *et al.* 2015) and given *P. martini* is also negatively affected by habitat development, it should be prioritized for conservation. In comparison, despite sufficient data, our prediction that *H. mabouia* will displace native *G. antillensis* and introduced *L. lugubris* is not supported by the LVM. The weakly negative correlation between *G. antillensis* and *H. mabouia* in the LVM may indicate displacement is just starting. Additionally, predation has been documented between *H. mabouia* and juveniles of other gecko species including *G. antillensis* (Dornburg *et al.* 2011, 2016) but predation is not well-represented by LVM correlation coefficients (Zurell *et al.* 2018) and may obscure the signal of competition. Although the predicted displacements of *G. antillensis* and *L. lugubris* are due to fitness differences only, examples of displacement due to fitness differences in reptiles are scant, despite fitness differences being implicated in invasion success in other systems (Godoy & Levine 2014; Gross *et al.* 2015; Ocampo-Ariza *et al.* 2018). Displacement due to fitness differences could manifest due to one species occupying limiting resources like diurnal refuges (e.g., Williams *et al.* 2016) or monopolizing temporal dimensions of resources (Rudolf 2019). Interestingly, our framework predicted the exclusion of *H. frenatus* by *H. mabouia* due to low niche differences. The LVM indicates this exclusion is not yet occurring, likely due to the recent arrival of *H. frenatus*, but the ecological similarity and low fitness differences between the two species suggest priority effects may dictate which species ultimately persists on Curaçao (Fukami 2015). This result is also pertinent for the Greater Caribbean region where introduced geckos including *H. mabouia* and *H. frenatus* are rapidly spreading (Behm *et al.* 2019; Perella & Behm 2020).

In conclusion, our framework can be applied in the early stages of an invasion to focus investigations of competitive displacement of native species. The framework helps provide theory-informed management of natural systems, facilitating the development of strategies for addressing the increasing threat that invasive species pose worldwide.

## ACKNOWLEDGEMENTS

We thank C.G. Irian and K.A. Langhans for help with field data collection; M. Vermeij, CARMABI, and numerous residents in Curaçao for permissions and permits; H.R. Assour and C.K.A. Lynch for morphological data collection; N. Belouard and T.M. Swartz for valuable comments on the manuscript, and P. M. Phillips for help with Fig. 2. This work was funded by Temple University and by Nederlandse Organisatie voor Wetenschappelijk Onderzoek, Grant Number: 858.14.041.

