## Supplementary Information for "Predicting and quantifying coexistence outcomes between resident and invading species using trait and abundance data"

Jocelyn E. Behm^1*^, Jacintha Ellers^2^, Wendy A. M. Jesse^2^, Tyler J. Tran^1^, and Matthew R.

Helmus^1^

^1^Integrative Ecology Lab, Center for Biodiversity, Department of Biology, Temple University, 1925 N 12^th^ Street, Philadelphia, PA 19122, USA

^2^Department of Ecological Science – Animal Ecology, Vrije Universiteit Amsterdam, Amsterdam, The Netherlands

*Corresponding author information
Jocelyn Behm

1925 N 12^th^ Street
Philadelphia, PA 19122


**Contents**

1. Supplemental Text S1

A) Supervised Landcover Classification Methods

B) Considerations on selecting LVM R-packages

2. Supplemental Results:

A) Survey Sample Sizes

Table S1 – Sample sizes per species for niche and fitness difference calculations

B) Niche Differences

Figure S1 – Correlations among α-niche morphological and microhabitat

variables

Table S2 – Loadings for α-niche traits in PCA

Figure S2 – Correlations among β-niche landcover variables

Table S3 – Loadings for β-niche traits in PCA

C) Fitness Differences

Table S4 – Effect of sampling date and habitat on proportion of gravid females

Table S5 – Effect of sampling date and habitat on proportion of juveniles

D) Coexistence Outcomes

Figure S3 – Coexistence space based on native-native species pairs

E) Joint Species Distribution Latent Variable Model Results

Table S6 – Likelihood ratio tests showing the improvement of fit for including

species-species correlation in LVMs for each buffer radius

Figure S4 – Effect of development on all six species (including HEFR) across all

buffer radius sizes

Figure S5 – Mean pairwise species correlations across species pair types

**SUPPORTING INFORMATION**

**1. Supplemental Text**

*A.) Supervised land cover classification for Curaçao*

Four Landsat 8 Collection 1 images were combined to create a nearly cloudless image mosaic of Curaçao (paths: 5-6 paths, rows: 51-52, dates: 24-12-2015, 28-7-2016, 4-8-2016, 26-12-2016). We classified images from single dates separately and performed extensive *posthoc* corrections on the classifications to minimize atmospheric and temporal effects (Song et al. 2001). We used the maximum likelihood supervised classification tool in ArcGIS 10.4 (ESRI, Redlands, CA, USA) to categorize pixels of the four images into eight categories: water, high-density vegetation, low-density vegetation, high-density development, low-density development, bare ground, cloud, and cloud shadow (see complete explanation and tutorial in Zhu 2016 pg. 269-272). We then combined the four classified Landsat images into a mosaic and scanned its entirety by eye with Google Earth imagery to correct all misclassified pixels.

*B.) Considerations on selecting latent variable model (LVM) R-packages*

Across ecological subfields, LVMs are gaining traction as useful analytical tools and as such there is an array of LVM R packages to choose from (e.g., Hui 2016, Niku et al. 2019, Tikhonov et al. 2020). For our system, the gllvm package (Niku et al. 2019) was best because it uses a maximum-likelihood approach that avoids the convergence issues Bayesian methods have with datasets with small numbers of species like ours (Hui, F.K.C. *pers. comm.*). Nonetheless, all LVM approaches where models are fit appropriately should provide qualitatively similar results. When selecting an LVM approach, we advocate testing several packages, if possible, to qualitatively compare results obtained with each package and ultimately select the package with the LVM that fits the data best and provides the best functionality to answer the research question at hand.

**2. Supplemental Results**

**A) Survey Sample Sizes**

We used data recorded from 288 geckos across 41 sites for our α-niche, β-niche and fitness calculations for predicting coexistence outcomes. For quantifying coexistence outcomes in progress, we used abundance data from 837 geckos across 26 sites (Table S1).

**Table S1**: Sample sizes per species for niche and fitness difference calculations and LVMs

| Species | *n* individuals for α-niche and fitness | *n* sites for β-niche | *n* sites for LVMs | *n* individuals for LVMs |
| --- | --- | --- | --- | --- |
| *Gonatodes antillensis* | 52 | 24 | 24 | 370 |
| *Phyllodactylus martini* | 83 | 35 | 19 | 188 |
| *Thecadactylus rapicauda* | 5 | 6 | 5 | 9 |
| *Hemidactylus frenatus* | 5 | 3 | 2 | 5 |
| *Hemidactylus mabouia* | 79 | 42 | 15 | 237 |
| *Lepidodactylus lugubris* | 12 | 10 | 5 | 28 |

**B) Niche Differences**

**Figure S1:** Correlations among α-niche morphological and microhabitat variables. Size of the circle and color intensity indicate strength of correlation; blue circles indicate positive correlations and red circles indicate negative correlations. Blank cells indicate the correlation was not statistically significant.

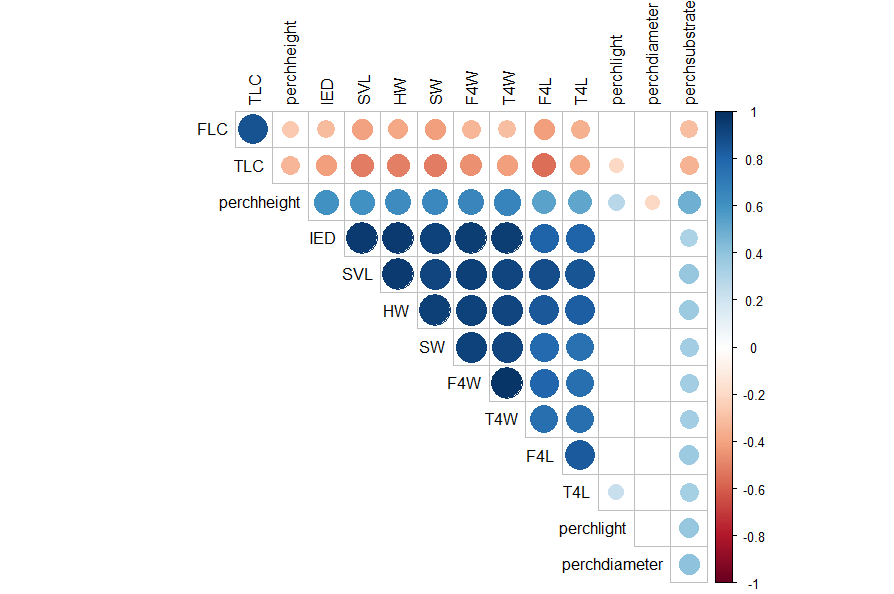

**Table S2:** Loadings for α-niche traits in PCA. Morphological traits are: snout-vent-length (SVL), snout width (SW), head width (HW), inter-eye-distance (IED), finger 4 width (F4W), finger 4 length (F4L), finger 4 lamellae count (FLC), toe 4 length (T4L), toe 4 width (T4W), and toe 4 lamellae count (TLC). Microhabitat traits are: perch height (cm) (continuous), perch diameter (cm) (continuous, flat perches like walls were assigned a maximum perch diameter), perch lighting (binary: light / dark), and perch substrate (binary: natural/manmade).

| **Trait** | **PC1α** | **PC2α** |
| --- | --- | --- |
| SVL | 1.96 | -0.17 |
| HW | 1.95 | -0.23 |
| IED | 1.91 | -0.43 |
| SW | 1.91 | -0.24 |
| F4W | 1.91 | -0.39 |
| F4L | 1.79 | 0.04 |
| FLC | -1.00 | -1.19 |
| T4W | 1.89 | -0.43 |
| T4L | 1.73 | -0.12 |
| TLC | -1.21 | -1.08 |
| Perch height | 1.43 | -0.07 |
| Perch light | 0.52 | 0.88 |
| Perch diameter | -0.12 | 1.44 |
| Perch substrate | 0.93 | 1.15 |

**Figure S2:** Correlations among β-niche morphological and microhabitat variables. Size of the circle and color intensity indicate strength of correlation; blue circles indicate positive correlations and red circles indicate negative correlations. Blank cells indicate the correlation was not statistically significant. Land cover “traits” are percent human development (‘devel’), high-density vegetation (remnant manchineel (*Hippomane mancinella*) forests; ‘hiveg’) and low-density vegetation (dry forests and scrub land; ‘lowveg’). Number after land cover trait indicates radius of buffer in m.

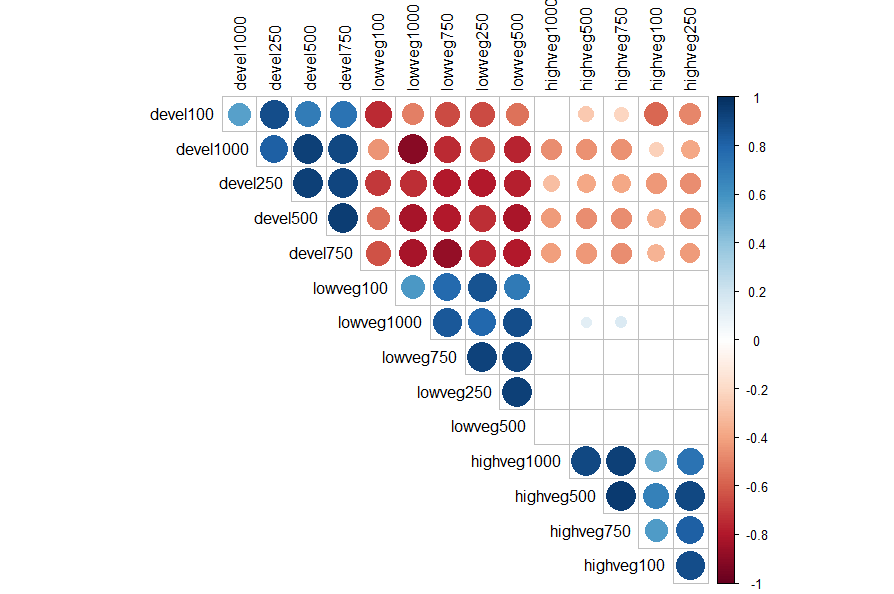

**Table S3:** Loadings for β-niche traits in PCA. Land cover “traits” are percent human development (‘devel’), high-density vegetation (remnant manchineel (*Hippomane mancinella*) forests; ‘hiveg’) and low-density vegetation (dry forests and scrub land; ‘lowveg’). Number after land cover trait indicates radius of buffer in m.

| **Trait** | **PC1β** | **PC2β** |
| --- | --- | --- |
| Devel100 | 1.67 | -0.05 |
| Devel250 | 2.01 | -0.05 |
| Devel500 | 2.02 | 0.07 |
| Devel750 | 2.04 | 0.00 |
| Devel1000 | 1.89 | 0.07 |
| Hiveg100 | -0.84 | -1.41 |
| Hiveg250 | -0.97 | -1.73 |
| Hiveg500 | -0.93 | -1.82 |
| Hiveg750 | -0.96 | -1.70 |
| Hiveg1000 | -0.85 | -1.67 |
| Lowveg100 | -1.43 | 1.03 |
| Lowveg250 | -1.67 | 1.14 |
| Lowveg500 | -1.72 | 1.01 |
| Lowveg750 | -1.82 | 0.79 |
| Lowveg1000 | -1.75 | 0.64 |

**C) Fitness Differences**

Results from linear models testing whether there was an effect of sampling date (Julian day) or habitat type (developed/undeveloped) on the proportion of gravid females and proportion of juveniles recorded during surveys. Significant effects of time or habitat would indicate the metric may not be appropriate for calculating fitness differences without additional considerations.

**Table S4:** Effect of sampling date (Julian day) and habitat type (developed/undeveloped) on proportion of gravid females

|  | Julian Day | | | Habitat type | | |
| --- | --- | --- | --- | --- | --- | --- |
|  | **Estimate** | ***P-*value** | **R-squared** | **Estimate** | ***P-*value** | **R-squared** |
| GOAN | -0.020 | 0.280 | 0.050 | -0.455 | 0.213 | 0.066 |
| HEFR | 0 | NA | NA | NA** | NA | NA |
| HEMA | -0.003 | 0.666 | 0.006 | 0.054 | 0.875 | 0.001 |
| LELU | 0.024 | 0.072 | 0.391 | 0.500 | 0.407 | 0.100 |
| PHMA | -0.019 | 0.324 | 0.034 | 1.616 | 1.000 | <0.001 |
| THRA* | NA | NA | NA | NA | NA | NA |

*Gravid females were not recorded for THRA
**HEFR was only found in developed sites

**Table S5:** Effect of sampling date (Julian day) and habitat type (developed/undeveloped) on proportion of juveniles

|  | Julian Day | | | Habitat type | | |
| --- | --- | --- | --- | --- | --- | --- |
|  | **Estimate** | ***P-*value** | **R-squared** | **Estimate** | ***P-*value** | **R-squared** |
| GOAN | 0.004 | 0.901 | 0.001 | 1.365 | 0.044 | 0.164 |
| HEFR | 0 | NA | NA | NA* | NA | NA |
| HEMA | -0.038 | 0.531 | 0.027 | 0.786 | 0.666 | 0.0123 |
| LELU | 0.022 | 0.454 | 0.146 | 0.800 | 0.432 | 0.160 |
| PHMA | -0.006 | 0.536 | 0.023 | -0.308 | 0.112 | 0.135 |
| THRA* | 0 | NA | NA | 0 | NA | NA |

*HEFR was only found in developed sites

**D) Coexistence Outcomes**

**Figure S3:** Pairwise niche and fitness differences for native-native species pairs along the four niche axes: PC1α (resource use axis), PC2α (microhabitat axis), PC1β (development axis), and PC2β (vegetation structure axis). The gray point clouds indicate the bootstrapped values and are bound within the 95% confidence ellipses (black line). Points are labeled with species pair names using the same abbreviations as in Fig. 2. The high niche and low fitness differences along PC1α indicate it should be used for defining coexistence space. Patterns of partitioning along the β−niche axes are used to confirm that the native species are interacting at spatial scales relevant for coexistence. The low niche differences along both β−niche axes indicate the native species are likely interacting at spatial scales where competitive exclusion can operate and supports the use of PC1α for calculating coexistence outcomes.

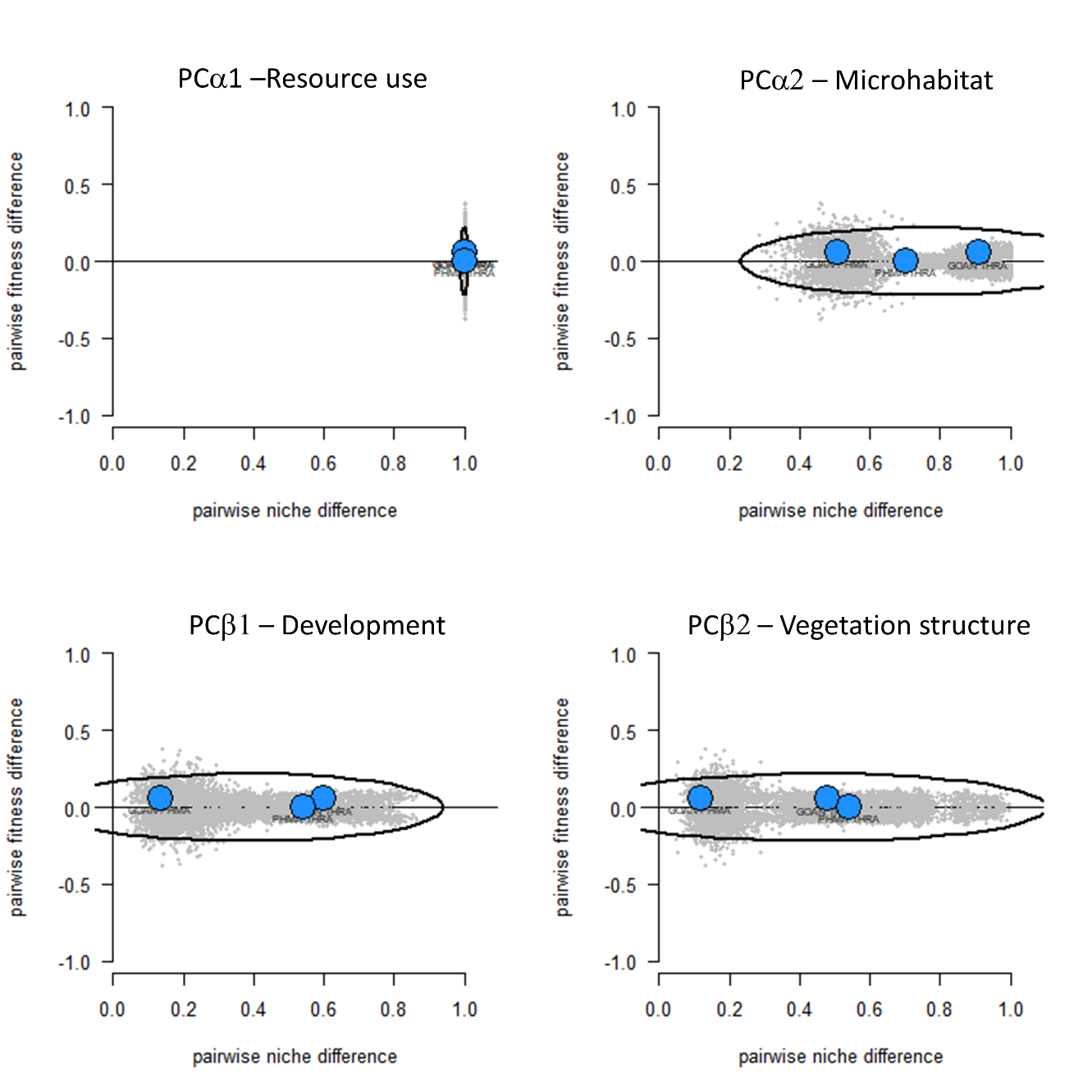

**E) Joint Species Distribution Latent Variable Model Results**

**Figure S4:** Effect of development on all six species from LVM. These are the same data plotted in Fig. 5 in the main text, but also includes *H. frenatus* (HEFR).

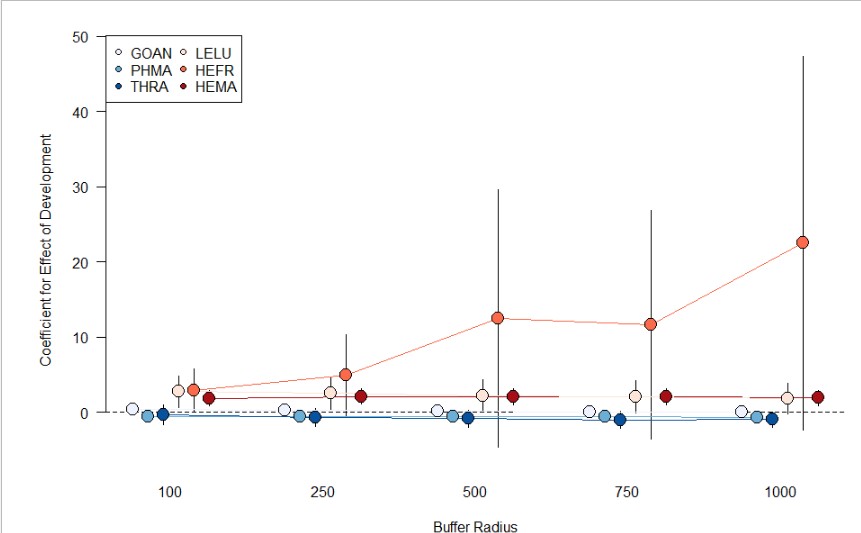

**Table S6:** Likelihood ratio tests showing improvement of fit for including species-species correlations in LVMs for each buffer radius

| **Buffer** | **Deviance** | **Df Difference** | ***P*-value** |
| --- | --- | --- | --- |
| 100m | 576.79 | 12 | <0.001 |
| 250m | 597.71 | 12 | <0.001 |
| 500m | 605.75 | 12 | <0.001 |
| 750m | 600.16 | 12 | <0.001 |
| 1000m | 607.05 | 12 | <0.001 |

**Figure S5:** Mean pairwise correlation coefficients +/- standard error for each species across all native and introduced species across the 5 buffer sizes. Points with no standard error lines have standard errors smaller than the point. Open points indicate correlation coefficients are not statistically different from zero. Point color indicates the type of species pairing. Species name abbreviations are the same as in Fig. 2.

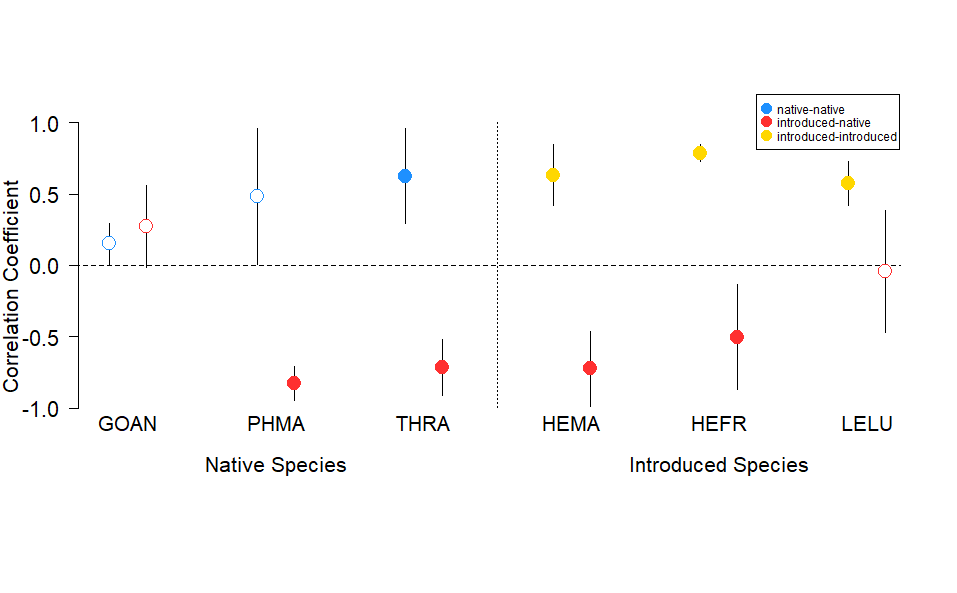
